# Spatiotemporal dynamics during niche remodeling by super-colonizing microbiota in the mammalian gut

**DOI:** 10.1101/2022.10.21.513299

**Authors:** Guillaume Urtecho, Thomas Moody, Yiming Huang, Ravi U. Sheth, Miles Richardson, Hélène C. Descamps, Andrew Kaufman, Opeyemi Lekan, Florencia Velez-Cortes, Yiming Qu, Lucas Cohen, Deirdre Ricaurte, Travis E. Gibson, Georg K. Gerber, Christoph A. Thaiss, Harris H. Wang

## Abstract

While fecal microbiota transplantation (FMT) has been shown to be effective in reversing gut dysbiosis, we lack an understanding for the fundamental processes underlying microbial engraftment in the mammalian gut. Here, we explored a murine gut colonization model leveraging natural inter-individual variations in gut microbiomes to elucidate the spatiotemporal dynamics of FMT. We identified a natural ‘super-donor’ consortium that universally engrafts into diverse recipients and resists reciprocal colonization. Temporal profiling of the gut microbiome showed an ordered succession of rapid engraftment by early colonizers within 72 hours followed by a slower emergence of late colonizers over 15-30 days. Moreover, engraftment was localized to distinct compartments of the gastrointestinal tract in a species-specific manner. Spatial metagenomic characterization suggested engraftment was mediated by simultaneous transfer of spatially co-localizing species from the super-donor consortia. These results offer a mechanism of super-donor colonization by which nutritional niches are expanded in a spatiotemporally- dependent manner.

## INTRODUCTION

The mammalian gut microbiome is composed of hundreds to thousands of bacterial species that co-exist symbiotically with their host and provide key metabolic and protective functions^1–3^. Despite being subjected to the harsh gastrointestinal (GI) environment and experiencing constant washout and nutritional shifts, the gut microbiome establishes reproducibly across individuals during early development and eventually reaches an equilibrium by adulthood^4^. Various environmental factors such as exposure to xenobiotics, antibiotics, or diet can lead to altered microbiome compositions and increased susceptibility to colonization by pathogens and pathobionts^5^. The recent success of fecal microbiota transplantation (FMT) to treat a disturbed gut microbiome suggests a robust process by which a healthy microbiome can be restored^6^. However, the detailed dynamics, mechanisms, and principles by which microbes successfully engraft into a resident community remain unclear.

The stability and malleability of a microbiome is shaped by various ecological properties including networks of inter-microbial interactions that manifest spatially and dynamically over time^7–9^. Metabolic interactions arise from commensal or mutualistic degradation of complex substrates that support multiple species in a consortium^10–12^. For example, *Bacteroides* in the gut are known to excrete various carbohydrate degradation enzymes that in concert liberate different sugars from dietary polysaccharides^10^. Similarly, diverse microbes spanning the length of the gut participate in the deconjugation and step-wise biotransformation of host-secreted bile acids, which alters local biochemical environments resulting in dramatic effects on gut biogeography^13^. Often, these metabolic activities reinforce positive-feedback loops that gradually result in systemic changes to the gut environment over sustained periods of time^14,15,16^. Mapping these interspecies interactions are key for assessing the stability of the microbiome and its susceptibility to colonization by other microbes.

The ability to colonize and shape microbial communities by introducing foreign microbiota is a quintessential goal of FMT therapies. Despite many successes, these therapies sometimes exhibit mixed outcomes that vary between different combinations of donors and recipient^17,18^. Curiously, “super donors” that consistently engraft in a variety of recipients have been reported^19^. While this phenomenon has been generally linked to species richness and diversity of donor communities, it is currently unclear what specific mechanisms or determinants are responsible^18,20–23^. The maturation of these therapies is thus reliant on developing an understanding of several key questions: Why do some strains engraft when others do not? Is variability in engraftment success due to donor or recipient composition or are there other factors? As the compositions of these microbial communities change, does their spatial structure change to resemble the donor? Answering these questions will require effective models of FMT and detailed dissection of the spatiotemporal dynamics and ecological interactions occurring within the gut microbiome.

In this work, we use a murine model system that exploits natural variations in the gut microbiome to study the temporal and spatial dynamics of microbial engraftment during FMT. This model recapitulates key features of human FMT including recipient heterogeneity, temporal successions, and diet dependencies while also providing tunable experimental parameters that are difficult to control in human FMT trials. We show that microbiomes of C57BL/6 mice acquired from different vendors exhibit variable outcomes when subject to pairwise FMT and identify a “super donor” consortium. Transplantation and successful engraftment of distinct taxa within this consortium occurred over short and long timescales. Further, we examine the spatial distribution of microbial engraftment across the recipient GI tract and identify key genetic factors associated with engraftment across different areas of the murine gut. Finally, spatial metagenomics was used to study how the micron-scale spatial structure of microbial communities is affected by engraftment, which revealed that spatial reorganization of the microbiota occurred concurrently with altered metabolic capacities of the FMT recipient microbiome. These results introduce a conceptual mechanism wherein colonizing microbes remodel metabolic niches temporally and spatially within gut environments to facilitate microbial colonization.

## RESULTS

### A natural murine model of gut microbiome variability and diversity

To develop a robust murine model for FMT dynamics studies, we first explored whether the murine gut microbiome exhibited the same degree of intra-host variability that is commonly observed in human populations^16,24^. Previous work suggested that genetically identical mice sourced from different commercial vendors had distinct gut microbiota^25–27^. To verify these findings, we obtained conventional C57BL/6 mice from four different suppliers (Jackson Labs, J; Taconic, T; Charles River, C; and Envigo, E) and performed 16S sequencing on their fecal matter (**Figure 1A**). Indeed, we observed that the gut microbiome from different suppliers had significantly distinct compositions in terms of taxa or OTUs (Operational Taxonomic Units) present (PERMANOVA, *p* = 0.001) and differences in alpha diversity (**Figure 1B, S1A**). Mice from Envigo displayed the greatest diversity, marked by high levels of *Prevotellaceae* and *Muribaculaceae*, while Taconic mice had the lowest diversity with an elevated proportion of *Firmicutes* relative to *Bacteroides* (**Figure S1B**). Importantly, the measured evenness of these mouse cohorts is within range of what is observed between healthy human cohorts and those with gastrointestinal disorders^28,29^ (**Figure S1C**). Therefore, the inter-vendor variability of the murine gut microbiome may be a tractable surrogate model for studying the principles guiding microbiota transfer between natural assemblages, which could help reveal shared properties underlying human FMT engraftment and outcomes.

**Figure 1.**
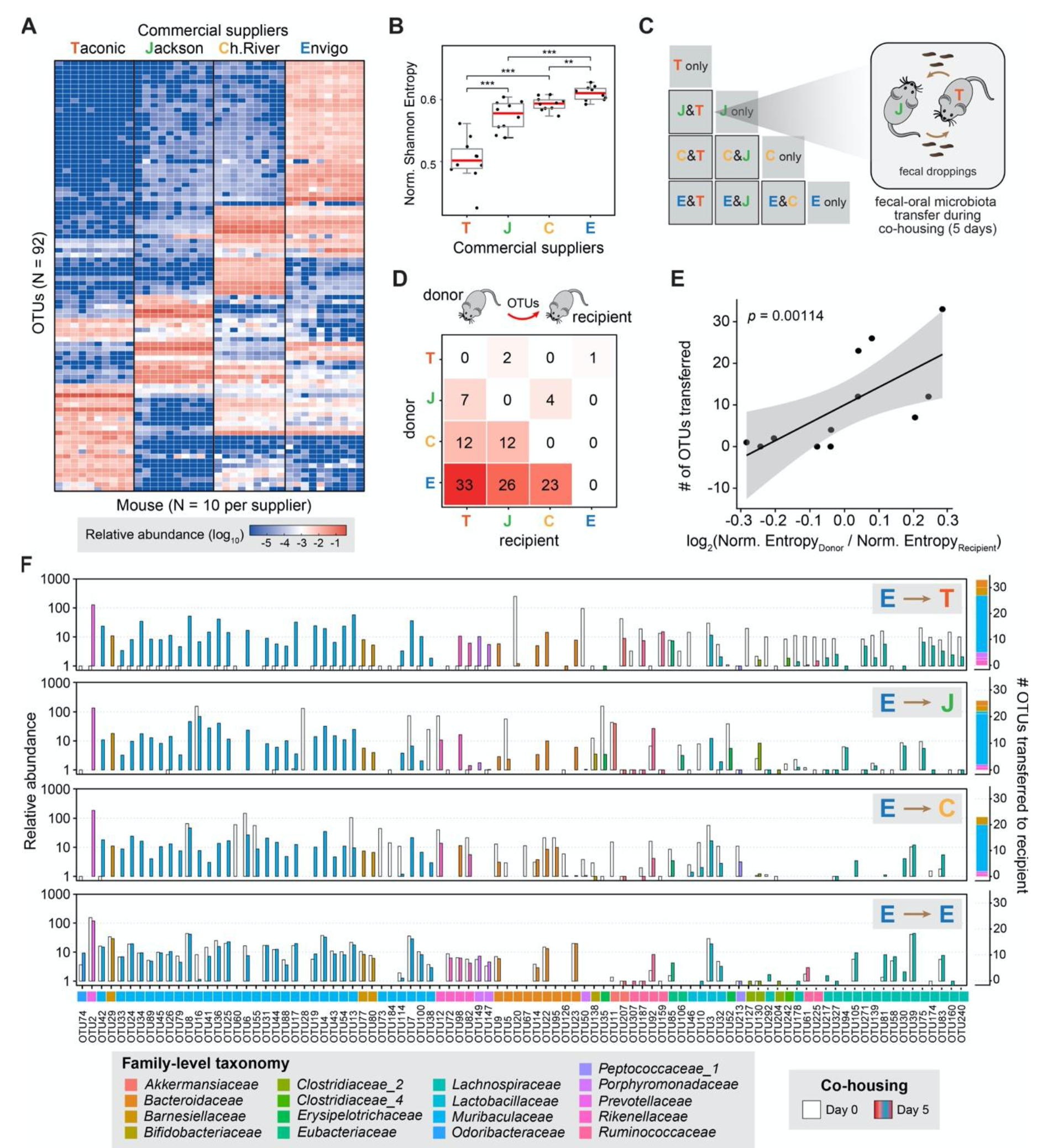
Diverse murine gut microbiomes exhibit variable outcomes to pairwise FMT (A) Microbiome composition of BL57BL/6 mice sourced from four vendors by 16S rRNA sequencing (N = 10 mice / vendor). **(B)** Shannon diversity index of mouse microbiomes (N = 10, Wilcoxon rank-sum test, *** = *p* < .001). **(C)** Pairwise fecal microbiota transfer (FMT) model by cohousing female C57BL/6 mice from different vendors. **(D)** Number of OTUs transferred between mice from different vendors. **(E)** The number of OTUs transferred from donor to recipient correlates with the ratio of their normalized Shannon Entropies. **(F)** (left) Relative abundance of OTUs at Day 0 and Day 5 of cohousing with Envigo mice. OTUs are ordered by Phylogenetic distance(right). Number and taxonomic identity of OTUs transferred from Envigo microbiome to various recipients.

### Variable engraftment of gut microbiota in a murine model of FMT

The inter-vendor variability in murine gut microbiomes is a useful property for an experimental model of microbial gut colonization that mimics the process of FMT in humans^30,31^. We therefore aimed to implement a simple, yet robust protocol that does not require pre- conditioning of the microbiome (e.g., use of antibiotics), and can leverage the natural variations in gut microbiomes of otherwise genetically identical mice. It is well established that cohoused mice from the same cage have nearly identical gut microbiomes because of fecal-oral transmission via coprophagy^30,32^. Leveraging this behavior, we cohoused mice from four different suppliers in a pairwise manner and profiled their gut microbiota by fecal 16S sequencing before (Day 0) and five days after (Day 5) cohousing (**Figure 1C**). Interestingly, we observed variable outcomes in terms of the number of OTUs transferred between different pairs of mice (**Figure 1D**). In most cases, bi-directional transfer of taxa occurred between different mice, with the number of unique OTUs transferred scaling linearly with the ratio of normalized entropy of the donor microbiota (**Figure 1E**). Members of the Envigo microbiome, which exhibited the greatest diversity, were capable of engraftment into all other microbiota and were highly resistant to reciprocal colonization. Strikingly, a group of 23-33 Envigo OTUs effectively transferred to all different recipient microbiota (**Figure S2**). Members of this “super-donor” consortium were predominately of the *Muribaculaceae* family but also included other taxa, including *Bacteroidaceae* and *Prevotellaceae spp.* (**Figure 1F**). While consistent sets of OTUs were transferred from Envigo to all recipients, recipient-specific transfers were also observed, including a pair of *Porphyromonadaceae* OTUs (OTU149 and OTU147) that were uniquely transferred to Taconic mice. These observations suggest a robust but complex ecological process underlying the observed colonization rather than random outcomes as predicted by the neutral theory of community assemblages^33,34^.

In addition to acquisition of new taxa, several OTUs initially present in the recipients were displaced after exposure to the Envigo microbiota. Often, this displacement coincided with transfer of phylogenetically related species from Envigo, which may indicate competition leading to replacement within similar ecological niches. For instance, multiple Charles River *Muribaculaceae spp.* (OTU60, OTU73, OTU184, OTU114) were displaced by ∼18 *Muribaculaceae spp.* transferred from Envigo (**Figure 1F**). Similarly, native Jackson and Taconic microbiomes contained high levels of single *Bacteroidaceae spp.* (OTU5 and OTU20, respectively) which were depleted concurrently with transfer of Envigo *Bacteroidaceae*, (OTU9, OTU14, OTU22, and OTU23). In other cases, native OTUs appear to be displaced in the absence of any phylogenetically similar taxa present from Envigo. Taconic mice contained a distinct population of 7 *Lachnospiraceae* OTUs (OTU139, OTU160, OTU174, OTU217, OTU240, OTU271, OTU327) that decreased in abundance (Wilcoxon rank sum exact test, *p* = 0.0006) yet no Envigo *Lachnospiraceae* were transferred. Therefore, Envigo microbiota appear to exhibit a “super-donor” phenotype that is sometimes observed in human FMT trials^22^. Recalcitrance of Envigo microbiota to invasion or displacement by other microbiota highlights this dominant persistence and colonization resistance phenotype.

### Ordered temporal microbiota transfer during murine FMT

To better elucidate the temporal dynamics of microbial transfer during our FMT model, we performed fecal 16S profiling of Jackson (Jax) mice cohoused with Envigo (Env) mice over 32 days. The Jax-Env pairing was chosen because they had significant differences in gut microbiome diversity (*p* = 0.00032, Wilcoxon rank sum test, **Figure 1B**). Beyond the dramatic changes to the Jax microbiome after five days of cohousing with Env mice, we were surprised to find that the microbiome of these recipient Jax mice (Jax^Env^) continued to change weeks after initiation of cohousing (**Figure 2A**). Microbes transferred to Jax^Env^ mice within the first five days were mostly *Lactobacillaceae* and *Muribaculaceae* whereas *Lachnospiraceae* emerged and reached moderate relative abundance over the latter half of this time-course experiment. Interestingly, Env *Lachnospiraceae* only began to colonize after 15 days, coinciding with the depletion of Jax *Lachnospiraceae*, which may suggest these incoming species are able to take advantage of a *Lachnospiraceae*-specific niche once it is vacated. Overall, differential abundance analysis showed 31 unique OTUs engrafted into Jax^Env^ mice, with 21 (67.7%) of these transferring within the first five days of cohousing (**Figures 2B, 2C**). At the conclusion of this time course, the Jax^Env^ microbiome exhibited higher population diversity and clustered more closely with the Env microbiome based on Principal Component Analysis (PCA) (**Figure S3A, S3B**). As controls, cohoused cage mates from the same vendor (i.e., Jax^WT^ or Env^WT^) did not lead to notable changes in the gut microbiome, nor did the recalcitrant Env mice when exposed to Jax microbiota (i.e., Env^Jax^) (**Figure S3C, Figure S3D**). These results show that transplanted microbiota emerge over both short (days) and long (weeks) time scales, which may indicate a gradual transition in the gut milieu towards stabilization.

**Figure 2.**
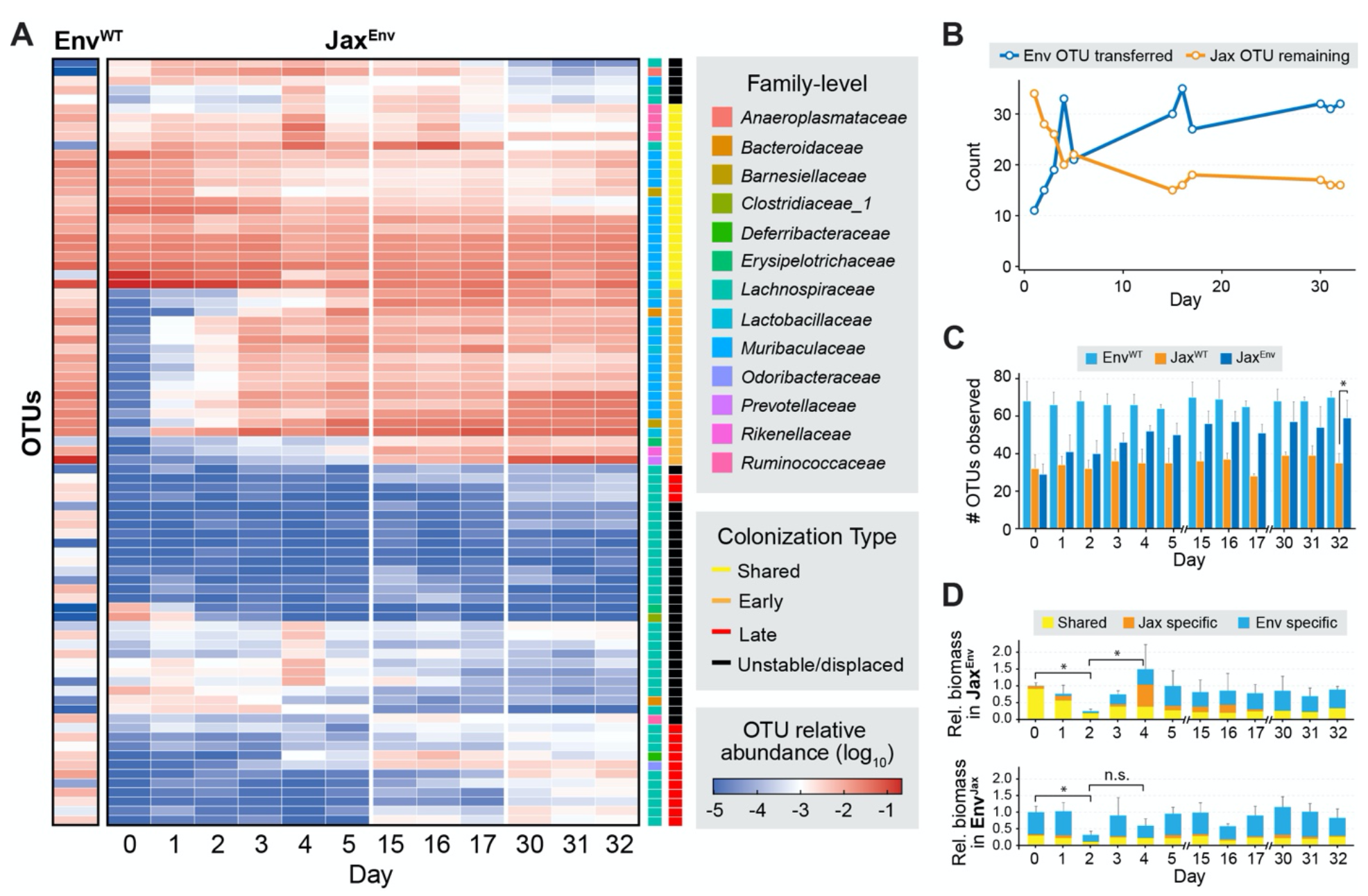
The transfer of microbes by FMT occurs over short and long-time scales. (A) Longitudinal 16S microbiome profiling of Jax^Env^ mice (N = 5). Detectable colonization by transferred OTUs occurs during both early (days 1-5) and late (days 15-32) sampling points. Representative Envigo microbiome is an average of day 1 Env-Env cohoused mice (N = 5). **(B)** Number of Env OTUs transferred and Jax OTUs remaining over longitudinal sampling. **(C)** Number of OTUs detected over time within each mouse cohort. We observed a significant increase in the number of OTUs present within Jax^Env^ mice compared to Jax^WT^ after 32 days of cohousing (Wilcoxon rank sum test, *p =* 0.029, n = 4). **(D)** Changes in bacterial biomass within feces of Jax^Env^ and Env^Jax^ mice. Biomass is colored by whether OTUs are uniquely found in microbiomes of Envigo donors (Donor specific), uniquely found in Jackson Recipients (Recipient specific), or observed in both. Values normalized to Day 0 of cohousing ( * = *p* < 0.05, one-sided Wilcoxon rank sum test).

Stable engraftment of new OTUs could correspond to expansion of niches in the gut, which would be reflected by an increase in carrying capacity of the community. We therefore assessed changes in bacterial density throughout FMT using absolute abundance measurements of fecal samples (Methods). Despite the consistent increase in unique OTUs observed over time, population load exhibited substantial temporal fluctuations. Overall biomass decreased over the first two days before a dramatic expansion followed by an equilibration (**Figure 2D**). By Day 30 the relative biomass corresponding to taxa specific to the Jax microbiota was entirely replaced by Env taxa in Jax^Env^ mice. While Env^Jax^ mice also experienced a bottleneck in population size at Day 2, there was no dramatic increase in biomass and the microbiome ultimately reverted to its original state. Neither control groups showed this phenomenon (**Figure S3E**). Our data therefore suggests that the convergence of microbial communities during FMT results from a transitionary state with a rapid and dramatic interval of population bottlenecking, followed by restructuring and re-equilibration of the new community.

### Microbial transplantation dynamics vary across murine gut compartments

The mammalian gut contains many ecological niches whose diverse environmental, biochemical, and ecological properties shape the gut biogeography, resulting in distinct microbial populations across different gut compartments^35^. Therefore, analysis of fecal pellets gives an incomplete picture of all changes occurring along the intestinal tract since fecal matter predominantly reflects the distal gut^36–38^. To explore engraftment dynamics across different compartments along the murine intestinal tract following FMT, we obtained GI samples upon necropsy at the conclusion of the 32-day Env-Jax cohousing experiment and performed 16S profiling of individual gut compartments spanning the entire intestinal tract (**Figure 3A**). PCA showed that the OTU composition of Jax^Env^ was more similar to that of Env across all gut compartments (**Figure S4A**). Interestingly, the taxonomic composition of the Jax microbiome appeared to be more uniform across gut compartments, whereas the gut microbiomes of Env^WT^ and Jax^Env^ cohorts were more stratified, with distinct microbial profiles in the small and large intestines. We confirmed this by performing an Analysis of Similarities^39^ (ANOSIM) and found that Jax gut compartments were significantly less dissimilar to each other than Env^WT^ or Jax^Env^ compartments (**Figure S4B**). Ultimately, FMT resulted in population remodeling across the entire length of the GI tract, shifting the 16S microbiome profiles of Jax^Env^ mice to resemble the Env microbiome across all gut compartments.

**Figure 3.**
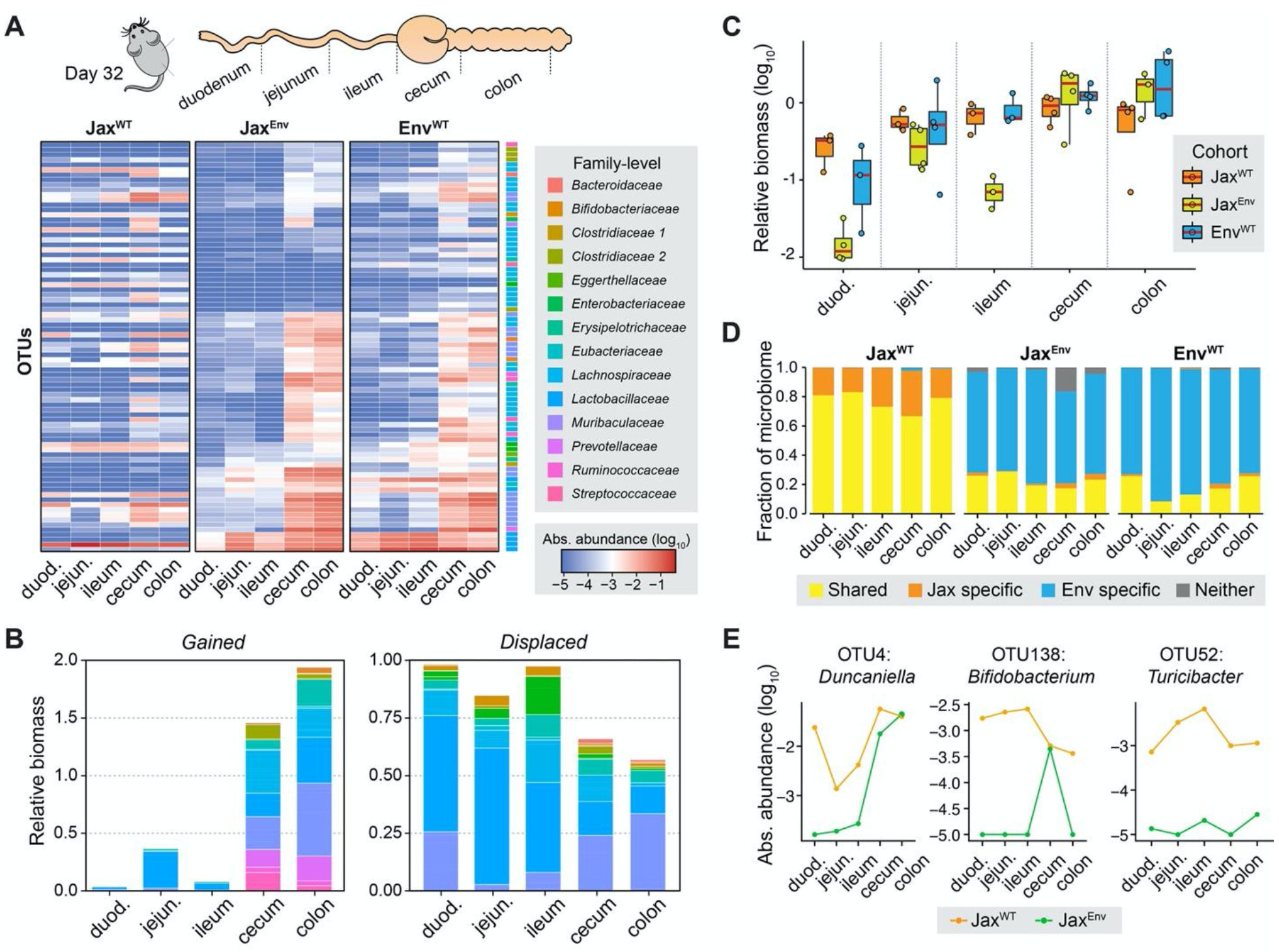
Microbial transplantation dynamics vary across murine gut compartments. (A) 16S profiling of luminal contents of mice cohorts after 32 days of cohousing. Rows are arranged by hierarchical clustering (Euclidean distance, Complete Linkage) of cohoused Jackson cohort. **(B)** Quantification of absolute bacterial biomass gained (left) and displaced (right) across all gut compartments stratified by taxonomic identity. Values are presented as relative to total Jax^WT^ biomass. **(C)** Absolute bacterial biomass in all gut compartments across mouse cohorts. **(D)** Proportion of bacterial biomass in each gut compartment uniquely found in Envigo donors (Env specific), uniquely found in Jackson Recipients (Jax specific), or observed in both. **(E)** Absolute abundance of select OTUs across gut compartments in Jax^WT^ and Jax^Env^ mouse cohorts.

To quantitatively assess how overall microbial communities were affected across different sections of the GI tract, we looked at changes in the biomass of different taxonomic groups in these areas. The composition of engrafting microbes varied dramatically across different compartments. Five species of Env-specific *Lactobacillaceae* were the primary colonizers of the small intestine, whereas diverse populations including *Muribaculaceae, Prevotellaceae, and Lactobacillaceae* colonized the cecum and colon of Jax^Env^ mice (**Figure 3B**). Conversely, a majority of the Jax recipient-specific biomass was displaced, especially within upper-GI compartments where overall bacterial biomass decreased by as much as 97.2% in the duodenum (**Figure 3C**). As was observed in our longitudinal profiling, OTUs specific to the Jax microbiome were nearly entirely replaced across all gut compartments while a proportion of the microbial taxa shared between the Env donor and Jax^Env^ decreased from 85-70% to 15-30% of the population of each gut compartment (**Figure 3D**). These data reflect the highly variable effects of FMT on microbial biomass and composition between intestinal compartments in the recipient.

Next, we explored whether the timing of ‘early’ or ‘late’ colonizing species was related to the areas they colonized in the gut. We compared the distribution of microbes to the order they colonized the GI tract observed in our longitudinal study (**Figure S4C**). Interestingly, “early colonizing” bacteria (mainly *Lactobacillaceae*) were more commonly observed in the upper GI whereas “late colonizers” were relatively enriched in the cecum and colon. Late-colonizing species nearly exclusively consisted of *Muribaculaceae* and *Lachnospiraceae*, the abundance of which are positively correlated with microbial production of deoxycholic acid (DCA)^40^. Considering conversion of primary bile acids to DCA is predominately facilitated by microbes in the upper GI^41^, early colonizing microbes may gradually alter DCA levels, enabling colonization by *Lachnospiraceae*. This raises the possibility that the late colonization phenomenon is due to early colonizers changing the biochemical properties of the gut before conditions are permissive to late colonizers.

In addition to engraftment of Env bacteria, the spatial distribution of many OTUs already found in Jax mice changed across the gut (**Figure 3E**). In some cases, these OTUs were consistently depleted across all gut compartments (OTU10, OTU52) whereas other microbes appeared to have been selectively depleted in specific gut compartments, but not others. The abundance of *Duncaniella* OTU4 decreased by 2-3 orders of magnitude in the small intestines yet remained unchanged in the colon. Similarly, OTU138, a *Bifidobacterium*, was found across all gut compartments in Jax mice, yet became restricted to the cecum in Jax^Env^ mice. This shows that microbial transfer by FMT results in non-uniform colonization across different areas of the gut and alters the biogeography of native species.

### FMT outcomes reflect micron-scale spatial structuring of OTUs within recipient and donor communities

The gut microbiome exhibits local spatial structuring that reflects complex inter-microbial interactions driven by mutualistic, commensal, and competitive processes^42^. Given the dramatic changes to the microbiome of Jax recipients following FMT from Env donors (i.e., Jax^Env^), we characterized the changes to the spatial structure of the microbiome using MaPS-seq, a sequencing-based method recently developed in our lab to obtain micron-scale species spatial co-association information^43^. We compared the microbial spatial co-association from Jax^Env^ with that of the Env^WT^ and the Jax^WT^ controls (N = 4 each) before and after 32 days of cohousing. In total, 20,992 unique MaPS-seq particles (20-40 µm in diameter) were profiled to reconstruct the spatial organization across these animal cohorts.

First, we sought to identify spatial co-associations between taxa pairs. A frequentist approach was used to simulate the co-occurrence of two OTUs within particles by generating the null distribution of co-occurrences for all pairs of OTUs, and then determining which microbes co-occurred within particles significantly more or less frequently than expected by chance^44^. This analysis evaluated spatial co-associations between 7,430 microbial pairs and identified 292 statistically significant co-associated OTU pairs from the Jax^WT^ microbiome (adjusted p < 0.05, randomization test) and 494 in Env^WT^ (**Figure S5**). Hierarchical clustering of these co-association pairs revealed distinct groupings. In Jax^WT^, spatial associations were predominately found amongst *Clostridia* species and separated into four distinct groups (**Figure 4A)**. Groups 1 and 4 formed highly connected within-group co-associations. Groups 2 and 3 had less within-group co-associations, except for a few strongly co-localized OTU pairs (e.g., Group 2: OTUs 79 and 56, Group 3: OTUs 148, 27, 5, and 99). Interestingly, we detected strong anti-associations between Groups 1 and 4; some Group 3 members also had negative association with members of Groups 1 and 4, suggesting spatial segregation across groups. In Env^WT^, we also observed four main groups of co-associated taxa (**Figure 4B**). Group 1 was distinctly dominated by *Clostridia* species and exhibited negative co-associations with *Bacteroidia* communities found in Groups 2 and 4. In contrast, Group 4 displayed the most class-level diversity, containing a mix of *Clostridia*, *Bacteroidia*, and *Bacilli*; this group had negative associations with Group 2, which was primarily composed of highly co-associated *Bacteroidia*. Lastly, Group 3 also consisted mainly of *Bacteroidia*, albeit with weaker overall interactions compared to other *Bacteroidia*-centric communities. These distinct spatial co-localizations suggest an organized community structure in the Env^WT^ and Jax^WT^ microbiomes of ecologically segregated *Clostridia* and *Bacteroidia* taxa.

**Figure 4.**
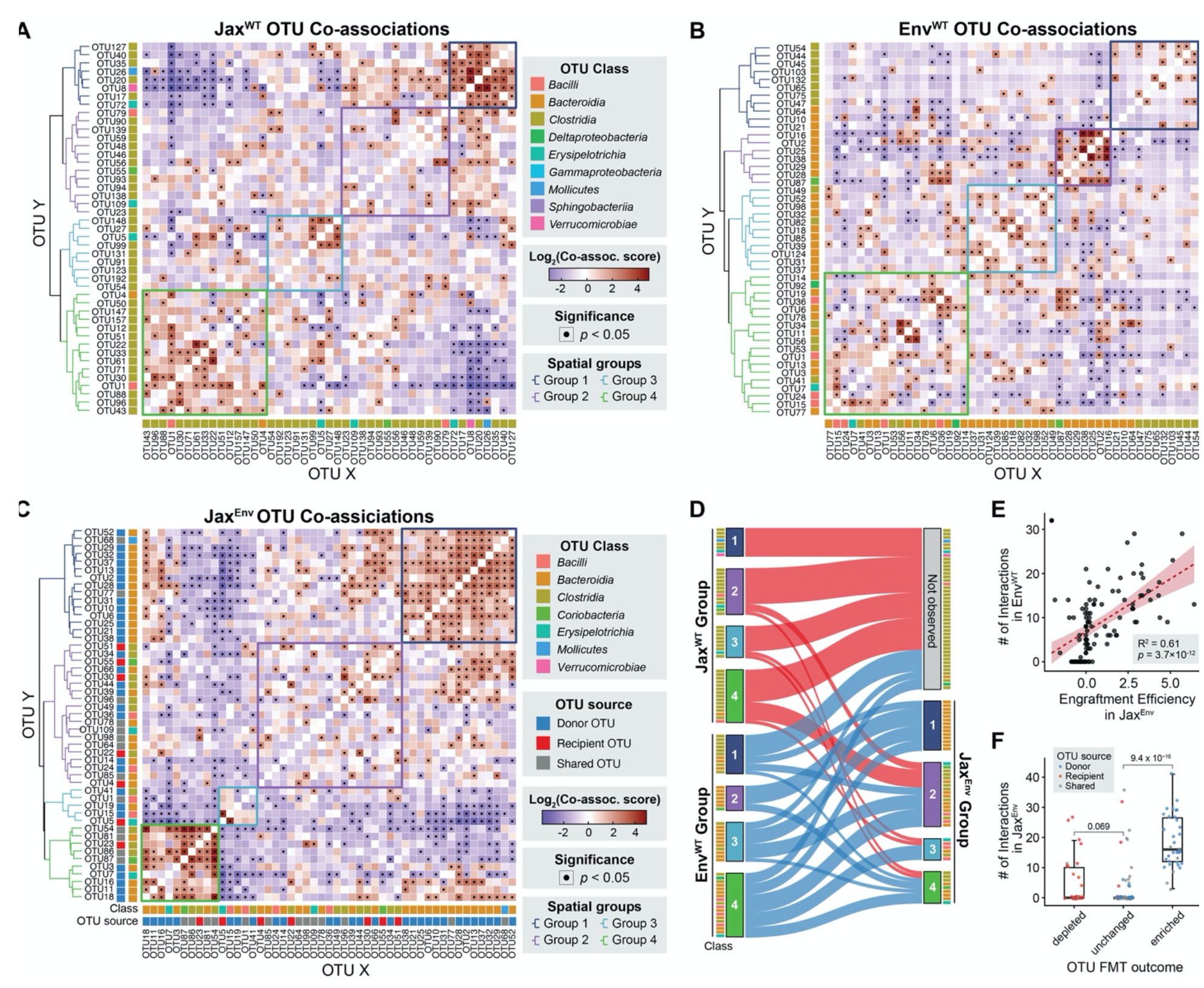
FMT results in dramatic restructuring of microbial spatial co-localizations. A-B) Co-association map indicating significant spatial co-associations amongst native OTUs unique to **A)** Env and **B)** Jax^WT^ mice. FMT outcomes determined by differential analysis comparing abundances in Jax^WT^ and Jax^Env^ mice. Rows and columns are clustered using Ward’s Linkage. **C)** Co-association map indicating significant spatial co-associations within the Jax^Env^ microbiota. OTUs are indicated depending on if they are uniquely found in Env, uniquely found in Jax, or shared. **D)** Sankey diagram indicating the transfer of OTUs from Jax/Env spatial subgroups to Jax^Env^ subgroups. **E)** Correlation between the number of observed spatial-associations in the Env microbiome and engraftment in Jax^Env^ across Env-enriched OTUs. Engraftment Efficiency indicates the log2 fold change in abundance comparing Jax^WT^ and Jax^Env^ mice. **F)** Number of interactions found amongst Env microbes separated by whether they were depleted, unchanged, or enriched in Jax^Env^ following cohousing (Wilcoxon rank sum test).

We then analyzed the Jax^Env^ microbiota to explore how spatial patterns can be altered following FMT (**Figure 4C**). In general, Jax^Env^ particles contained significantly more distinct OTUs per particle than Jax^WT^ particles (Jax^Env^: 3.76, Jax^WT^: 2.88, *p* < 2 x 10^-16^, Wilcoxon rank-sum test), indicating Env microbiota transfer into to the Jax community moderately increased species richness at the micrometer scale (**Figure S6**). We detected 499 significant spatial co-associations in Jax^Env^ that clustered into four groups (**Figure 4C**, **Figure S5**). While the Jax^Env^ microbiota was mostly comprised of Env OTUs, numerous *Clostridia* Jax OTUs were retained. Fascinatingly, some transferred Env OTUs reassembled into a spatial structure resembling their same configuration in the original Env^WT^ microbiota whereas others formed hybrid communities with Jax taxa. In Group 1, *Bacteroidia* species reunited within the recipient microbiome creating a community exclusively consisting of Env OTUs. This group exhibited intriguing relational dynamics: it had positive associations with Group 2, but a mixture of strongly negative and positive interactions with Group 4. Overall, Group 2 exhibited relatively weak spatial structuring aside from a strongly positive co-association between OTU30 and OTU96. Interestingly, this association was also observed in the Jax^WT^, and was preserved through FMT. Group 3 was driven by a particularly strong co-associations between an Env *Bacilli* (OTU15) and Jax *Erysipelotrichia* (OTU5) and these taxa were negatively associated with nearly all other members of the microbiome. Intriguingly, in Env^WT^, OTU5 formed strong spatial association with another *Erysipelotrichia* (OTU7) (**Figure 4B**). Not only did OTU15 form a strong association with the Jax OTU5, but it also became negatively associated with its Env^WT^ partner, OTU7. This observation highlights the dynamic ecological strategies microbes can employ to successfully adapt and colonize in the context of FMT.

We then explored whether microbes transferred by FMT retained their spatial groups through the transfer from the original microbiomes to Jax^Env^ (**Figure 4D**). Flow analysis demonstrated that while a fraction of each spatial group was preserved through transplantation, donor microbes predominantly formed novel subcommunities within the recipient gut environment. Spatial groups appeared to cluster more by their taxonomic composition than by their original group, indicating that taxonomic identity plays a more decisive role in the formation and stabilization of new microbial communities following transplantation. Jax^Env^ Group 1 was a hybrid community of *Bacteroidia* derived from multiple Env^WT^ groups, particularly those from Env Groups 2 and 3. Jax^Env^ Group 2 received the largest share of Jax taxa, most of which were co-associated before FMT in the Jax^WT^ Group 4. Intriguingly, all constituents of Env Group 2, which exhibited the most robust associations in Env^WT^ , achieved successful transfer.

Considering this observation, we examined whether the presence of spatial associations in the donor microbiome was predictive of FMT outcomes in the recipient. Indeed, the number of spatial associations by Env microbes in their Env^WT^ native microbiome was correlated to their engraftment success in the Jax^Env^ recipient (R^2^ = 0.61, *p* = 3.7 x 10^-12^) (**Figure 4E**). Amongst Jax taxa, microbes with the most interactions in Jax^WT^ were not necessarily more stable post-FMT. Rather, Jax microbes that remained stable following FMT were found to increase in the number of associations they exhibited in Jax^Env^ and these new associations were formed with Env taxa (**Figures S7**). Furthermore, bacteria significantly enriched in Jax^Env^ compared to Jax^WT^ had notably more spatial associations than microbes with either unchanged abundance (*p* = 9.4 x 10^- 15^, Wilcoxon rank sum test) or those depleted after FMT (*p* = 1.5 x 10^-7^, Wilcoxon rank sum test) (**Figure 4F**). Altogether, these results suggest microbes with the capacity to form spatially organized communities are better able to engraft compared to species that do not.

### Exploitation of open nutritional niches as a key determinant of FMT engraftment success

We hypothesized that the successful engraftment of spatially associated communities is facilitated by enhanced metabolic capabilities, enabled by synergistic mutualistic interactions such as cross-feeding. To catalog the metabolic diversity across mice gut microbiomes from different suppliers, we performed shotgun metagenomic sequencing on their fecal DNA, which yielded a 240 Gigabase (Gb) dataset that assembled into 457 metagenome-assembled genomes (MAGs) with annotated gene functions (Methods). Rather than relying on existing mouse strain databases, we performed *de novo* assembly to avoid database biases favoring different vendors or culturable strains. This collection of MAGs covers over 80% of all genus-level diversity across the four distinct C57BL/6 gut microbiomes (**Fig 5A**). Upon taxonomically assigning MAGs, we confirmed that Envigo consortia harbored a higher number of *Muribaculaceae*, 35 distinct MAGs, which are known to be prolific mucin foragers with diverse polysaccharide degradative capacities^29^. More detailed genomic characterization (Methods) of the Carbohydrate-Active Enzyme (CAZyme) repertoire within *Muribaculaceae* MAGs revealed that these microbes contained a set of unique CAZyme families (GH148, GH155, GH158, GH121, GH116, GH47), which may indicate these bacteria are able to utilize a broader range of dietary polysaccharides (**Figure 5B**, **Methods**). Beyond CAZyme differences, we investigated whether certain KEGG pathways were enriched between these metagenomes but did not find any significant differences (**Figure S8A**). These findings highlight that the mouse gut microbiomes derived from different vendors exhibit distinct microbial profiles and variability in CAZyme composition, which may lead to differences in polysaccharide utilization between these cohorts.

**Figure 5.**
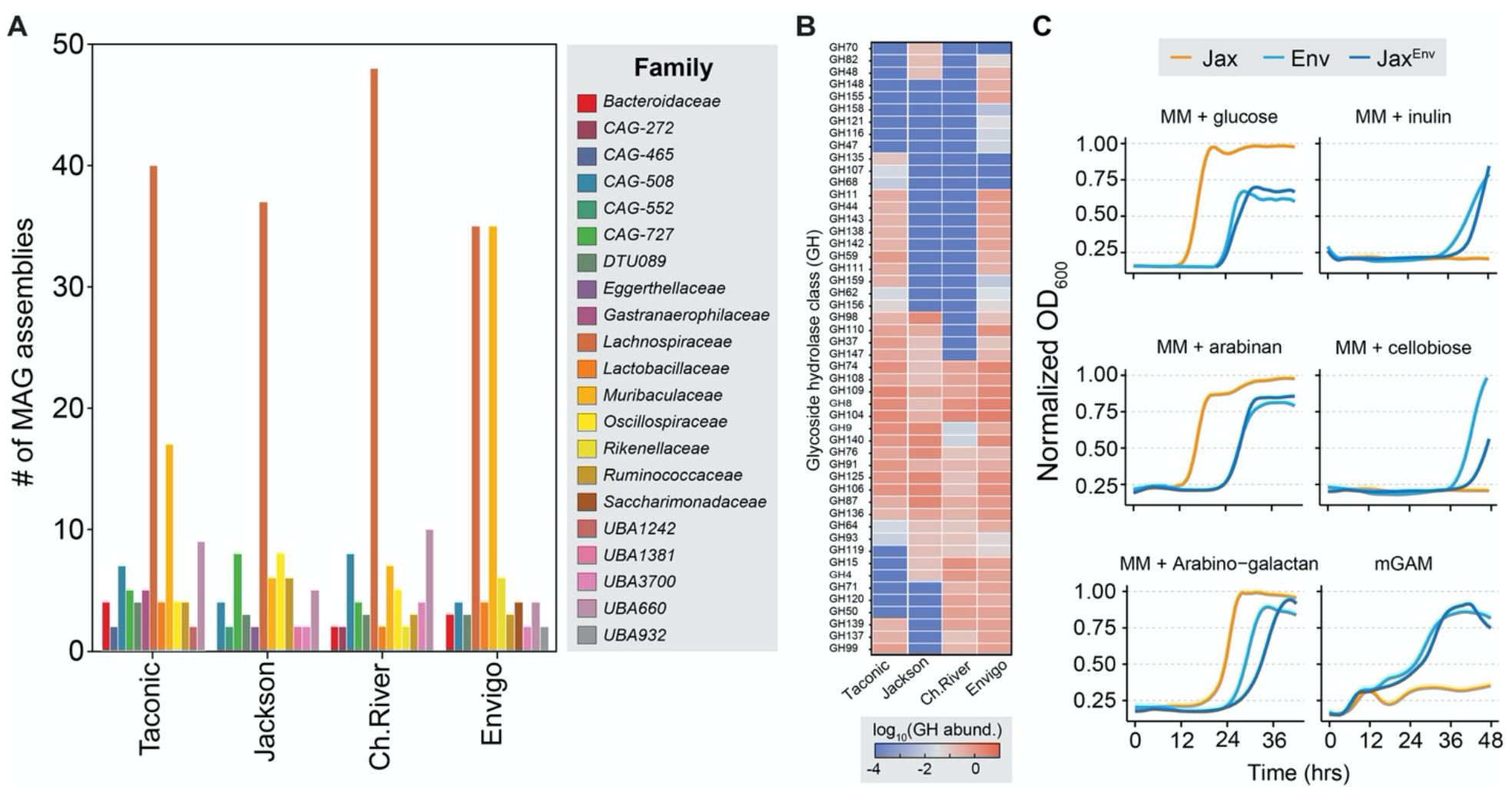
Envigo microbiota harness unique glycoside hydrolases to metabolize diverse carbohydrate substrates. (A) Number of metagenome-assembled genomes (MAGs) associated with each vendor and their family-level taxonomic distribution. **(B)** Abundance of glycoside hydrolase genes within *Muribaculaceae* MAGs from each vendor. **(C)** OD600 Growth assays of fecal communities acquired from Jax^WT^, Env^WT^, and Jax^Env^ mice after five days of cohousing. Communities were inoculated in defined minimal media supplemented with single sources of carbohydrates (indicated).

Given that the most prolific colonizers in the Env microbiota consisted of spatially organized *Bacteroidia* communities, we sought to explore how Jax and Env *Bacteroidales* communities differed in their metabolic capacities to access nutrient niches in the gut environment. *Bacteroidales* are the primary metabolizers of complex dietary polysaccharides in the gut and work in concert to break down these macromolecules into consumable subunits^10^. Interestingly, gut colonization of engineered probiotics can be enhanced by fiber supplementation, in cases where the probiotic is the sole species capable of metabolizing that fiber^45,46^. Therefore, we hypothesized that the super-colonizing phenotype of Env *Bacteroidia* is driven by their ability to metabolize previously inaccessible dietary fibers within the Jax recipient GI tract.

We evaluated the abilities of fecal communities from Env^WT^, Jax^WT^, and Jax^Env^ mice to utilize various complex polysaccharides. Growth assays on fecal communities was performed over 48 hours using defined minimal media supplemented with a panel of simple and complex carbohydrates to characterize the range of accessible polysaccharides. Growth is only possible if the community can break down the supplemented complex polysaccharides. We observed striking differences in growth profiles between the microbiomes. Env^WT^ and Jax^Env^ microbiota could utilize certain complex dietary polysaccharides (e.g., inulin, cellobiose) as the sole carbon source while Jax^WT^ microbiota could not (**Figure 5C**). Moreover, the complex modified gifu anaerobic media (mGAM) could support growth of Env^WT^ and Jax^Env^ but not Jax^WT^ microbiota. Conversely, the native Jax^WT^ microbiota grew faster in glucose, arabinan, and arabinogalactan indicating specialization for using these resources. 16S profiling of saturated communities showed that more diverse *Bacteroidales*-enriched populations grew from the Env^WT^ and Jax^Env^ communities in all conditions (**Figure S8B**). This is consistent with the idea that mixed communities of engrafted *Bacteroidalia* work cooperatively to break down previously unused dietary polysaccharides into available carbohydrates. This data therefore suggest that the Env microbiome can access a broader set of carbohydrate nutrients to supplement their growth, which allow them to exploit unfilled nutrient niches in the recipient Jax gut microbiota. FMT from Env microbiota imparts the ability for this ecosystem to metabolize additional carbohydrates, which may expand the accessible nutrient niches within the recipient gut and increase the carrying capacity of the environment.

### Using humanized mouse gut microbiomes to simulate human FMT outcomes

Finally, we explored the use of our murine model to simulate FMT dynamics between humans. Notably, the humanized mouse model allows the study of reciprocal FMT between two microbiomes which is not practically feasible in human FMT studies. We acquired fecal samples from three representative human donors of the three major human gut microbiome enterotypes^47^. These individuals, H1, H3, and H5, were dominated by *Ruminococcae* (enterotype 3), *Prevotellaceae* (enterotype 2), and *Bacteroidaceae* (enterotype 1), respectively. We gavaged germ-free mice with these fecal samples and observed engraftment after nine days that only partially resembled their respective human donors (**Figure S9A**). Humanized mice were predominately colonized by families *Bacteroidaceae* and *Akkermansiaceae* regardless of the enterotype of the donor, whereas *Prevotellaceae* and *Ruminococcaceae* were poorly represented. Moreover, the microbiomes of humanized mice were similar at the family-level but varied greatly when comparing OTU-level resolution and diversity metrics (**Figure S9B**). Although humanized mice microbiomes did not fully recapitulate the microbiomes of their donors^48^, PCA showed the greatest similarity to their respective donors (**Figure S9C**). We then performed pairwise co-housing between humanized mice to explore FMT outcomes between human microbiomes (**Figure 6A**). After nine days of cohousing, fecal 16S sequencing revealed two main groups of microbes that exhibited similar transfer dynamics and were generally enriched for members of the order *Clostridiales* (**Figure 6B**). PCA showed that when M5 mice were cohoused with M1 or M3 mice (i.e. M5^M1^, M5^M3^) the M5 microbiome composition shifted to resemble the other microbiomes (**Figure 6C**). On the other hand, cohoused M1^M3^ & M3^M1^ mice form a new grouping resembling an intermediate between M1^WT^ and M3^WT^. Examining the number of OTUs transferred, we found M1 and M3 mice transferred 30 and 26 species to M5 mice, and these OTUs were predominately *Clostridiales* (**Figure 6D**). The large number of *Clostridiales* transferred to M5 mice is notable given that the M5 mouse microbiome contained the fewest number of *Clostridiales* OTUs (**Figure S9D**). Therefore, this may indicate the M5 microbiomes contained a vacancy in *Clostridiales* niches that were exploited to promote engraftment of additional *Clostridia* and that the taxonomic ‘completeness’ of a microbiome may determine permissiveness to engraftment. This is consistent with the results of a recent meta-analysis of FMTs that showed community-dissimilarity between donor and recipient was a strong predictor of engraftment^23^. Overall, these results show that humanized murine gut models produce interesting FMT outcomes that could provide further insights into the determinants of microbiota transfer and colonization in humans.

**Figure 6.**
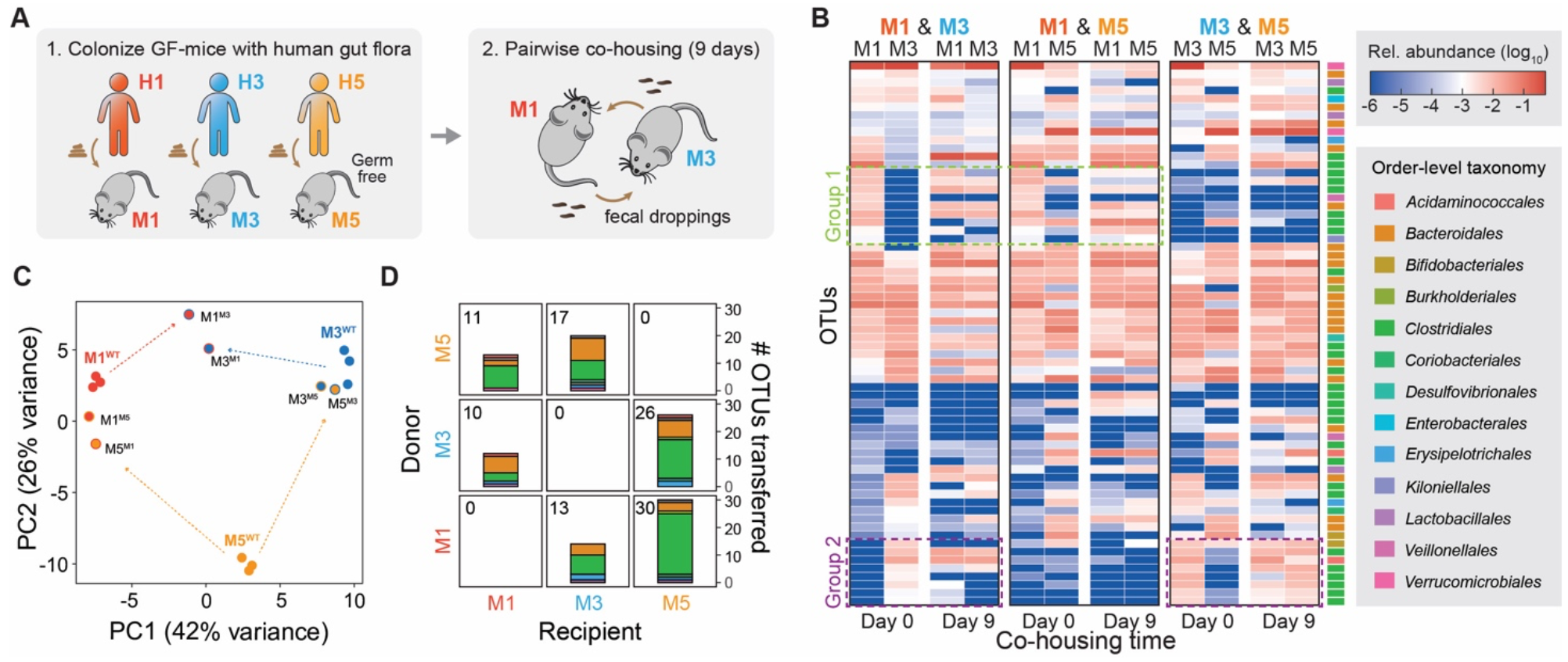
Humanized mouse microbiomes simulate human FMT outcomes. **(A)** Pairwise fecal microbiota (FMT) transfer model of gnotobiotic female C57BL/6 mice harboring ‘humanized’ microbiomes from individuals spanning the three canonical enterotypes. **(B)** 16S profiling of mouse fecal communities following nine days of cohousing separated by pairs (N=2 per pair). Clusters 1 & 2 indicate observed microbial transfer events. Rows arranged by hierarchical clustering. **(C)** PCA of euclidean distance in OTU composition comparing humanized mice before and after cohousing. The primary label indicates the original mouse microbiome and superscript indicates the cohousing partner. **(D)** Taxonomic composition of transferred OTUs between C57BL/6 mice harboring ‘humanized’ microbiomes. The number in the top left indicates the total number of OTUs transferred. OTU color scheme matches **(B)**.

## DISCUSSION

In this study, we characterized microbial gut colonization in a murine FMT model that exploited natural microbiota variations in different mouse cohorts. Through pairwise co-housing and FMT experiments, we identified a “super donor” microbiome from Envigo mice suppliers capable of dominantly engrafting into other murine microbiomes and reciprocally resisting colonization by all other microbiomes. Longitudinal profiling of a Env-Jax FMT pair revealed kinetics of colonization and adaptation of the microbiome in Jax^Env^ groups, characterized by immediate and gradual engraftment and multiple surges in microbiota abundances. Spatially characterizing the microbiota across different GI compartment showed that Env and Jax^Env^ mice had similarly more segregated microbial populations across the GI tract than compared to Jax mice. Application of MaPS-seq to these microbiome samples showed these microbiomes contain spatially separated communities that transfer collectively but reorganize in the recipient during FMT. Finally, a humanized gut microbiome FMT experiment demonstrated the utility of this murine model for studying FMT potential between human cohorts.

Our observations highlight several key features underlying bacterial engraftment in the gut. Through kinetic studies, we found both short (days) and long (weeks) timescale emergence of transferred strains. We hypothesize that following initial FMT, microbes begin a gradual process of shaping the gut environment such that native taxa are suppressed, which ultimately paves the way for other transferred species to gain a foothold. This may occur due to changing of the biochemical properties or by the suppression of native species. Thus, the temporal dynamics and succession observed during FMT may mirror those seen in other examples of microbial colonization, such as following antibiotics or development^49–51[cite]^. Future studies should examine the metabolic and biochemical changes that occur during this process, which may reveal new facets about how microbial communities interact and are established. A particularly interesting observation from this temporal analysis was the sharp drop in overall microbial population load in the short-term following FMT followed by a bloom and subsequent stabilization. This observation may be explained by the results of a previous longitudinal study, which showed that dramatic microbial community transitions are often preceded by an initial population-level bottleneck before dense, stable communities are established^52^. This may be a general phenomenon that occurs during the merging of microbial communities and warrants further investigation to learn the inter-microbial dynamics responsible, which may include direct antagonism or the collapse of cross-feeding networks.

Using MaPS-seq, we showed for the first time the micron-scale consequences of FMT on communities within a recipient gut. This analysis revealed a key observation that microbes form spatially associated communities in the gut and these communities are reassembled in the recipient during FMT. Furthermore, these units may actively displace other native community units found in the recipient. Generally speaking, microbes exhibiting the greatest number of spatial associations in the donor microbiome were more successful at colonizing the recipient GI tract than those that do not. Given that spatial relationships may represent underlying ecological interactions, such as mutualism, cooperation, or competition, these spatial associations may provide crucial information for understanding the inter-species mechanisms affecting how microbial communities are established during FMT.

In this study, a positively co-associating group of *Bacteroidia* was found to transfer from Envigo donors to Jax recipients. Further functional studies in defined media showed that the Env microbiota are capable of broadly utilizing polysaccharides that are otherwise inaccessible to the Jax microbiota. While Jax microbes were capable of metabolizing fewer carbohydrates, they grew faster than Env microbes when provided appropriate resources, suggesting more of a specialist than generalist lifestyle. We speculate that the trade-off of generalist versus specialist communities may be an important factor in determining the success of FMT therapies and that generalist communities may be better suited for engraftment into recipients. Given that the mammalian intestinal tract is a dynamic environment, with constantly fluctuating resources and host-derived inputs^53,54^, generalists may be more effective at enduring these changing conditions and ultimately supplanting populations of specialists. Indeed, generalist communities have been found to outperform specialists during the merging of aquatic microbial communities under dynamically changing environmental conditions^55^. Another explanation for the success of generalist communities is that the broad range of nutrients they can utilize equips them to exploit unused nutritional niches within recipient gut environments as metabolically independent units^56^. The creation of nutritional niches through dietary supplementation has shown to be an effective tool for enabling engraftment of probiotic microbes^45,57^. However, our research reveals that unused niches in the recipient gut environment may be exploited to promote the successful transplantation of microbiota into the mammalian gastrointestinal tract.

This work systematically explored the spatiotemporal dynamics of microbial colonization following FMT. Future applications of this approach could better delineate the role of host-factors in FMT outcomes and under clinically relevant settings such as exposures to antibiotics and xenobiotics. Ultimately, we expect more detailed spatial, temporal, genomic, and metabolic characterizations of the gut microbiome FMT kinetics will lead to more predictive FMT models that can unlock the true potential of this microbiome manipulation approach for a variety of clinical applications.

## RESOURCE AVAILABILITY

### Lead contact

Further information and requests for resources and reagents should be directed to and will be fulfilled by the lead contact, Harris Wang.

### Materials availability

This paper does not report original materials.

### Data and code availability

● Raw sequencing data is available through SRA under BioProject ID: PRJNA1028308.
● Original code and processed datasets are available through https://github.com/gurtecho/Urtecho_et_al_FMT
● Any additional information required to reanalyze the data reported in this paper is available from the lead contact upon request.

## EXPERIMENTAL MODEL AND SUBJECT DETAILS

### Mouse lines

C57BL6/J Mice were separately purchased from Jackson Laboratory, Taconic Biosciences, Envigo, and Charles River Laboratories.

## METHOD DETAILS

### Animal procedures

6- to 8-week-old female C57BL6/J mice were obtained from different suppliers and allowed to acclimate to the animal facility for a week in cages of four mice. After one week, the bedding was exchanged between cages of mice from the same vendor to normalize their microbiota. To enable FMT by cohousing, after normalization, two mice from each cohort were transferred to a new cage along with two mice from a second cohort. In control groups, all cohoused mice were from the same vendor. Mice were fed Teklad global 18% protein (2018S).

### Mice feces collection and microbial DNA extraction

Fresh mouse fecal pellets were collected and kept on dry ice before being weighed and transferring to a -80°C freezer for long-term storage. Whole pellets were suspended in 1 mL PBS and mechanically separated using an inoculating loop. Genomic DNA (gDNA) of fecal microbiota were extracted using a silica bead beating-based protocol adapted from Qiagen MagAttract PowerMicrobiome DNA/RNA Kit [Qiagen 27500-4-EP], detailed fully in Ref^58^. For experiments in which absolute abundance was determined, 1 uL of saturated *Sporosarcina pasteurii* (ATCC 11859) culture was added to the sample prior to bead beating.

### Luminal Content Collection

The luminal contents of mice were extracted for 16S and metagenomic sequencing. Mice were euthanized and their intestinal tracts were dissected in a sterile hood. The small intestines were separated into three sections of equal length, and their gut contents as well as those of the cecum and large intestines were extruded into 1.5 mL tubes and transferred to dry ice using forceps. Samples were weighed and processed following the microbial DNA extraction protocol described above.

### 16S rRNA amplicon sequencing

16S sequencing of the V4 region for mice gut microbiota was performed using a custom library preparation and sequencing protocol with dual indexing strategy^58^. Briefly, a 20uL 16S-V4 PCR reaction was set up (1ng extracted gDNA; 1uL forward barcoded P5 primer; 1uL reverse barcoded P7 primer; 10uL NEBNext® Ultra™ II Q5® Master Mix [NEB M0544X]; SYBR Green I at 0.2x final concentration) and subjected to a quantitative amplification on a thermal cycler (98°C 30s; cycles: 98°C 10s, 55°C 20s, 65°C 60s; 65°C 5min; 4°C infinite). PCR reaction was stopped during exponential phase to avoid amplification bias (typically 13-16 cycles) and the cycling was skipped to the final extension step. Next, 16S-V4 amplicon libraries were pooled based on the fluorescence increase at the last cycle and subjected to gel electrophoresis. DNA bands at ∼390bp were excised from gel and purified using Wizard™ SV Gel and PCR Cleanup System (Promega A9282) following the manufacturer’s instructions. Purified libraries were sequenced on Illumina MiSeq platform (reagent kits: v2 300-cycles, paired-end mode) at 8 pM loading concentration with 25% PhiX spike-in (Illumina FC-110-3001). Custom sequencing primers were spiked into reagent cartridge (well 12: 16SV4_read1, well 13: 16SV4_index1, well 14: 16SV4_read2) following the manufacturer’s instructions.

### 16S rRNA amplicon analysis and OTU clustering

Raw sequencing reads of 16SV4 amplicon were analyzed by USEARCH v11.0.667^59^. Specifically, paired-end reads were merged using “-fastq_mergepairs” mode with default setting. Merged reads were then subjected to quality filtering using “-fastq_filter” mode with the option “-fastq_maxee 1.0 -fastq_minlen 240” to only keep reads with less than 1 expected error base and greater than 240bp. Remaining reads were deduplicated (-fastx_uniques) and clustered into OTUs (-unoise3) at 100% identity, and merged reads were then searched against OTU sequences (-otutab) to generate OTU count tables. Taxonomy of OTUs were assigned using the Ribosomal Database Project classifier trained with 16S rRNA training set 16.

### OTU Filtering and Count Normalization

OTU count tables were normalized to relative or absolute abundance and filtered by relative abundance for downstream analyses as follows. For experiments lacking spike-in controls, reads were normalized by relative abundance within each sample and OTUs with a relative abundance below 0.5% (averaged across biological replicates) were removed. Absolute abundance measurements were determined by normalizing relative abundance of all OTUs to spike-in OTU counts as well as the weight of the fecal pellet.

### Shotgun metagenome sequencing

Library preparation of shotgun metagenome sequencing was performed using the same gDNA used for 16SV4 amplicon sequencing. Briefly, Nextera libraries were prepared following a scale-down Tn5 tagmentation-based library preparation protocol with 2ng gDNA as input^60^. Libraries were sequenced on Illumina Nextseq 500/550 platform (2 x 75bp) and HiSeq platform (2 × 150bp) following the manufacturer’s instructions.

### Metagenome assembly and binning

Raw reads of shotgun metagenome sequencing were processed by Cutadaptv2.1^61,62^ with the following parameters “--minimum-length 25:25 -u 5 -U 5-q 15 --max-n 0 --pair-filter=any” to remove Nextera adapters and low-quality bases. To obtain metagenome-assembled genomes (MAGs), processed raw reads of each mouse cohort were first assembled using metaSPAdes v3.11.1^63^ with default parameters. Yielding contigs of each cohort were split into 10kb fragments to denoise assemby artifacts and then subjected to binning by MaxBin v2.2.6^63,64^, MetaBAT v2.12.1^65^, CONCOCT v1.0.0^65,66^, and MyCC^67^ (no version info) with default settings. Results from different tools were further integrated and corrected by DAS Tool v.1.1.1^68^ to generate a first round of metagenome bins. Raw reads were then aligned to metagenome bins using Bowtie2 v2.3.4^69^ in “—very-sensitive” mode and partitioned into bins based on alignments. Next, partitioned reads of each bin were assembled separately by Unicycler v.0.4.4^70^ with default setting to generate final MAGs. All MAGs were then evaluated for quality and contamination by Quast v4.6.3^71^ and CheckM v1.0.13^72^ and subsequently subjected to taxonomy annotation by GTDB-Tk v1.7.0^72,73^.

### Functional annotation of MAGs

Protein sequences of MAGs were annotated by Prokka v1.12^74^ with default settings and were used for functional annotation to assign CAZyme and KEGG terms. Briefly, reference protein sequences with specific CAZyme annotation or KEGG Orthology terms were downloaded from CAZyme database or KEGG database respectively, and homolog search was performed for MAG protein sequences against reference sequences using BLASTP v2.9.0+ with maximum targets no more than 50. BLAST targets with e-value < 0.0001 were considered as hits, and the CAZyme or KEGG Orthology terms annotation was then assigned to MAGs’ protein sequences based on their BLAST hits.

### MaPS-Seq Sample Collection

Fresh fecal pellets were collected and immediately transferred to tubes containing methacarn (60% methanol, 30% chloroform, 10% acetic acid). After 24 hrs of fixation, samples were transferred to 70% ethanol and stored at 4°C until use. Samples were processed following the MaPS-Seq protocol^43^. After fracturing and barcoding, 20-40 micron particles were isolated by size-exclusion filtering for sequencing. For each mouse, two technical replicates of approximately 20,000 particles were used for sequencing. Samples were sequenced on an Illumina NextSeq550 (2 x 250 bp).

### MaPS-Seq Particle Clustering

particles were clustered using the louvain algorithm, as implemented in the Seurat R package function FindClusters() with the resolution parameter set to 0.5.

### Spatial association analysis within MaPS-Seq Particles

A frequentist analysis was performed to identify spatial associations between OTUs. Briefly, OTU counts in each particle were binarized to create a matrix representing the presence or absence of each OTU in each particle. To simulate a null model of co-occurrence, we used the EcoSimR package v0.1.0 to randomly shuffle presence and absence counts and count the number of particles each OTU pair was found together for. This was performed 1000 times for each sample to generate a distribution of co-occurrence frequencies for each OTU pair. We then determined where the observed co-occurrence frequency laid along this distribution and calculated the corresponding Z-score and two-tailed P-value. P-values were adjusted using false-discovery rate and an adjusted p-value < 0.05 was considered significant. Network analysis was performed in R using the packages ggraph v2.0 and igraph v1.3.1 using co-occurrence Z-scores to indicate the magnitude of relationships.

### Polysaccharide Utilization Growth Assays

Fecal communities were grown in Bacteroides minimal media^75^ cultures supplemented with various polysaccharides. Fecal pellets were mechanically separated in 1 mL PBS and diluted 1:10 in PBS before being inoculated in a 96-well culture for a total dilution rate of 1:400. Cultures were supplemented with 10 mg/mL of a single carbohydrate. The resulting inoculated cultures were grown over 48 hrs in a Biotek Powerwave XS plate reader (product code: B-PWXS) taking OD600 measurements every 15 minutes and the data was exported for analysis in R.

### Ethical review

This study was approved and conducted under Columbia University Medical Center Institutional Animal Care and Use Committee (Protocol #AC-AABD4551) and complied with all relevant regulations.

## Acknowledgements

We thank members of the Wang laboratory for advice and comments on the manuscript. H.H.W. acknowledges funding support from the NSF (MCB-2025515), NIH (1R01AI132403, 1R01DK118044, 1R21AI146817), ONR (N00014-18-1-2237, N00014-17-1-2353), Burroughs Wellcome Fund (1016691), Irma T. Hirschl Trust, and Schaefer Research Award. G.U. was supported by the HHMI Hanna H. Gray Postdoctoral Fellowship (GT15182). T.M. is supported by NIH Medical Scientist Training Program (T32GM007367). R.U.S. was supported by a Fannie and John Hertz Foundation Fellowship and an NSF Graduate Research Fellowship (DGE-1644869).

M.R. and F.V.C. are supported by NSF Graduate Research Fellowships (DGE-1644869). G.K.G. acknowledges funding support from the NSF (MCB-2025515), NIH (1R01GM130777), and BWH President’s Scholar Award.

---

**Supplementary Figure 1.**
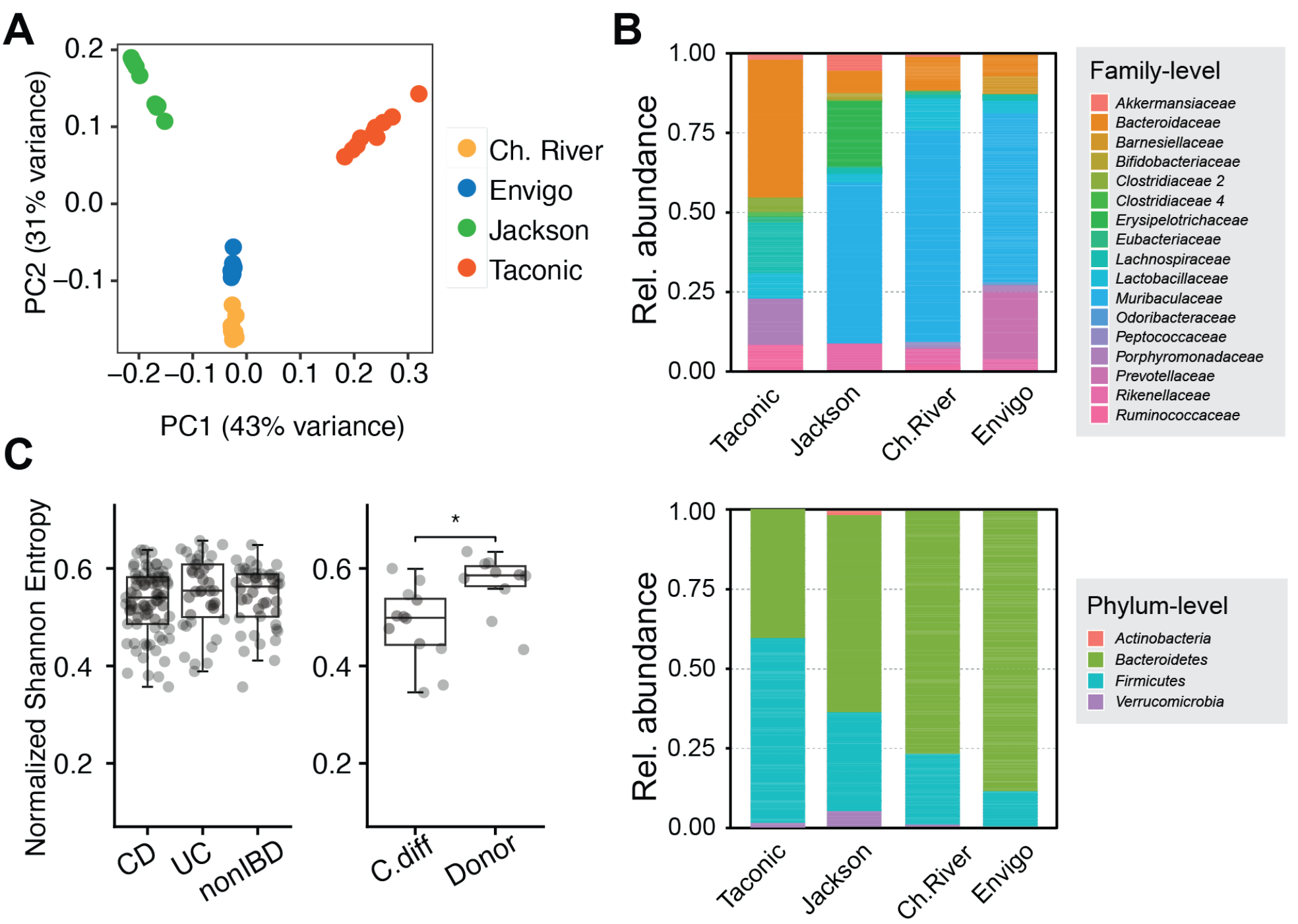
Microbiome OTU composition and functional annotations. (A) PCA of unweighted UniFrac distance comparing mouse 16S profiles from different vendors **(B)** Family-level (top) and Phylum-level (bottom) 16S composition of microbiomes from various vendors. **(C)** Shannon Diversity estimations of human fecal 16S profiles from healthy (non-IBD, Donor) and diseased (*p* = 0.01377, Wilcoxon rank sum test) (UC: Ulcerative Colitis, CD: Crohn’s Disease, C.diff: *Clostridioides difficile*) cohorts. IBD – Irritable Bowel Disease; UC – Ulcerative Colitis; CD – *Clostridioides difficile*.

**Supplementary Figure 2.**
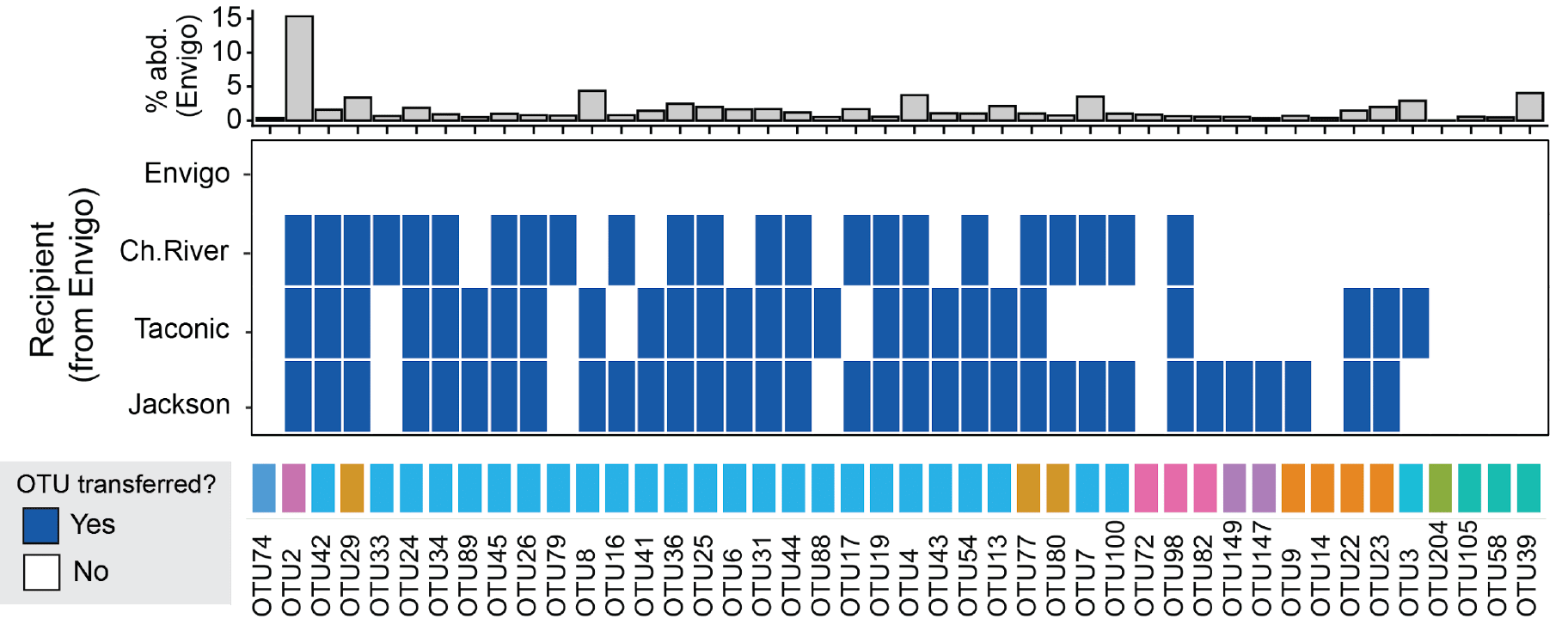
OTUs gained or displaced by cohabitation with Envigo. Summary of OTUs transferred from Envigo to various recipients via cohousing. (top) % abundance of each OTU in Envigo controls.

**Supplementary Figure 3.**
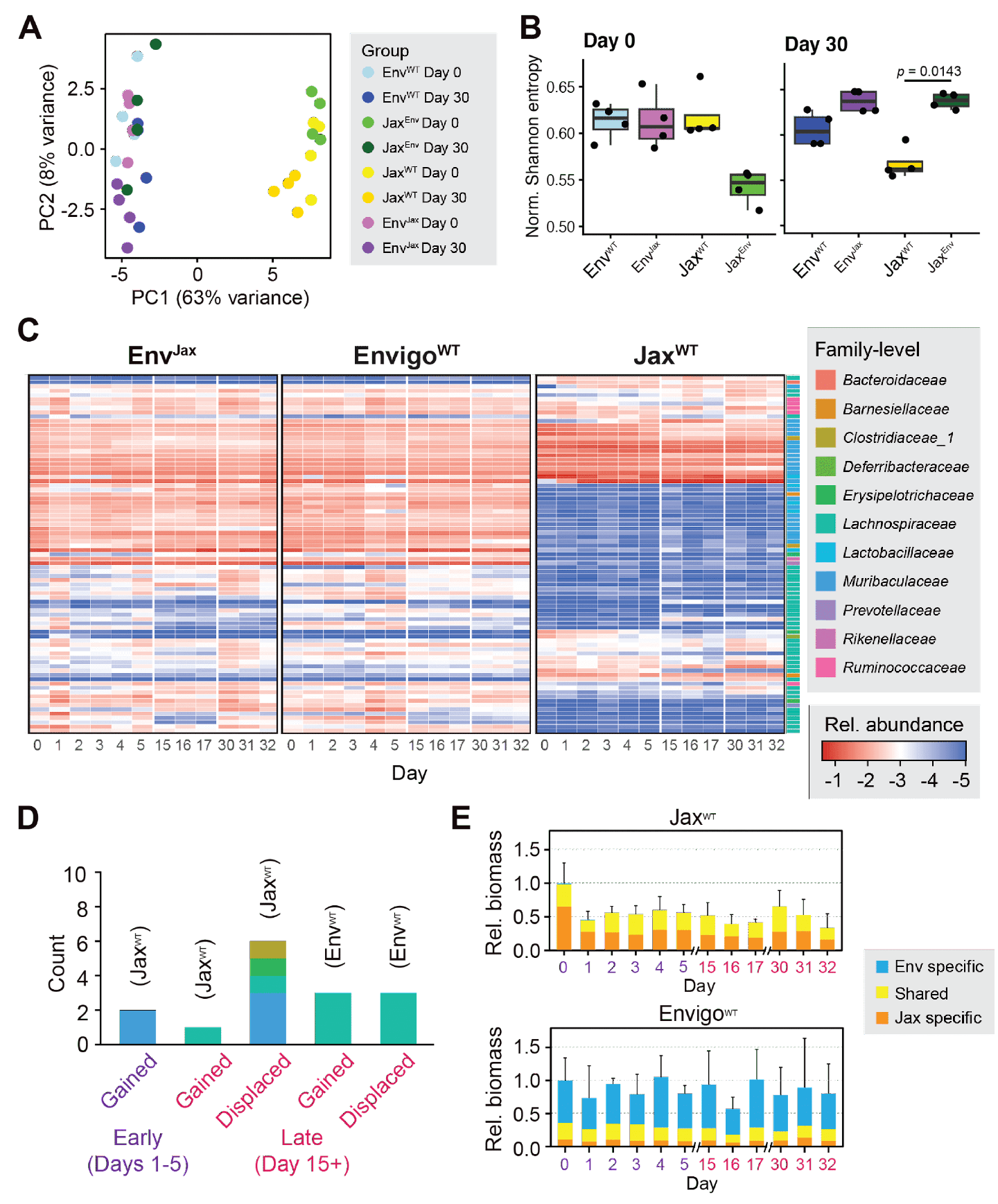
**(A)** PCA of 16S fecal microbiome composition between mouse cohorts at days 0 and day 30 of cohousing. **(B)** Shannon Diversity Index (SDI) of cohoused mice at day 0 and day 30 (N = 4). **(C)** Longitudinal 16S microbiome profiling of cohoused mice controls. (left) Envigo mice cohoused with Jackson mice, (middle) Envigo mice cohoused with other Envigo mice, (right) Jackson mice cohoused with other Jackson mice (N = 4). **(D)** OTU composition of microbes gained and lost during cohousing in non-mixed Jax^WT^ mouse controls and Env^WT^ mouse controls. **(E)** Relative bacterial biomass within feces of (top) Jax^WT^ mice non-mixed controls and (bottom) Env^WT^ mouse non-mixed controls. Biomass is colored by whether OTUs are uniquely found in microbiomes of Envigo donors (Env specific), uniquely found in Jackson Recipients (Jax specific), or observed in both (Shared).

**Supplementary Figure 4.**
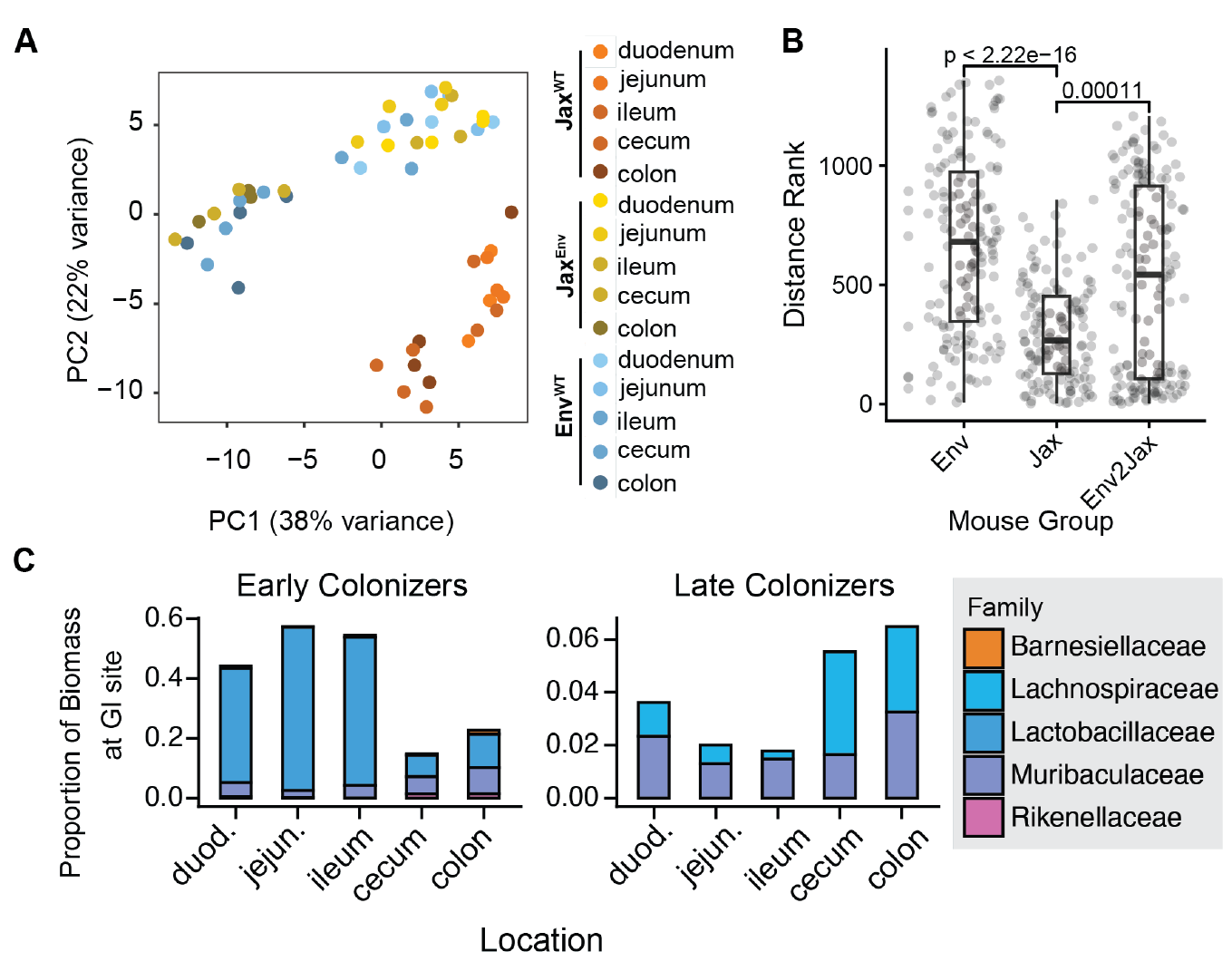
**A)** PCA of 16S microbiome composition across different gut compartments (N = 4). Jax^Env^ mice have greater variance between compartments and resemble the Env^WT^**. B)** Analysis of Similarities (ANOSIM) measurement of dissimilarity within mouse groups (Wilcoxon rank sum test). **C)** Relative proportion of Jax^Env^ mouse microbiome composed of (left) Early and (right) late colonizing OTUs at each GI site after cohousing. Sampling done at day 32 of mouse cohousing.

**Supplementary Figure 5.**
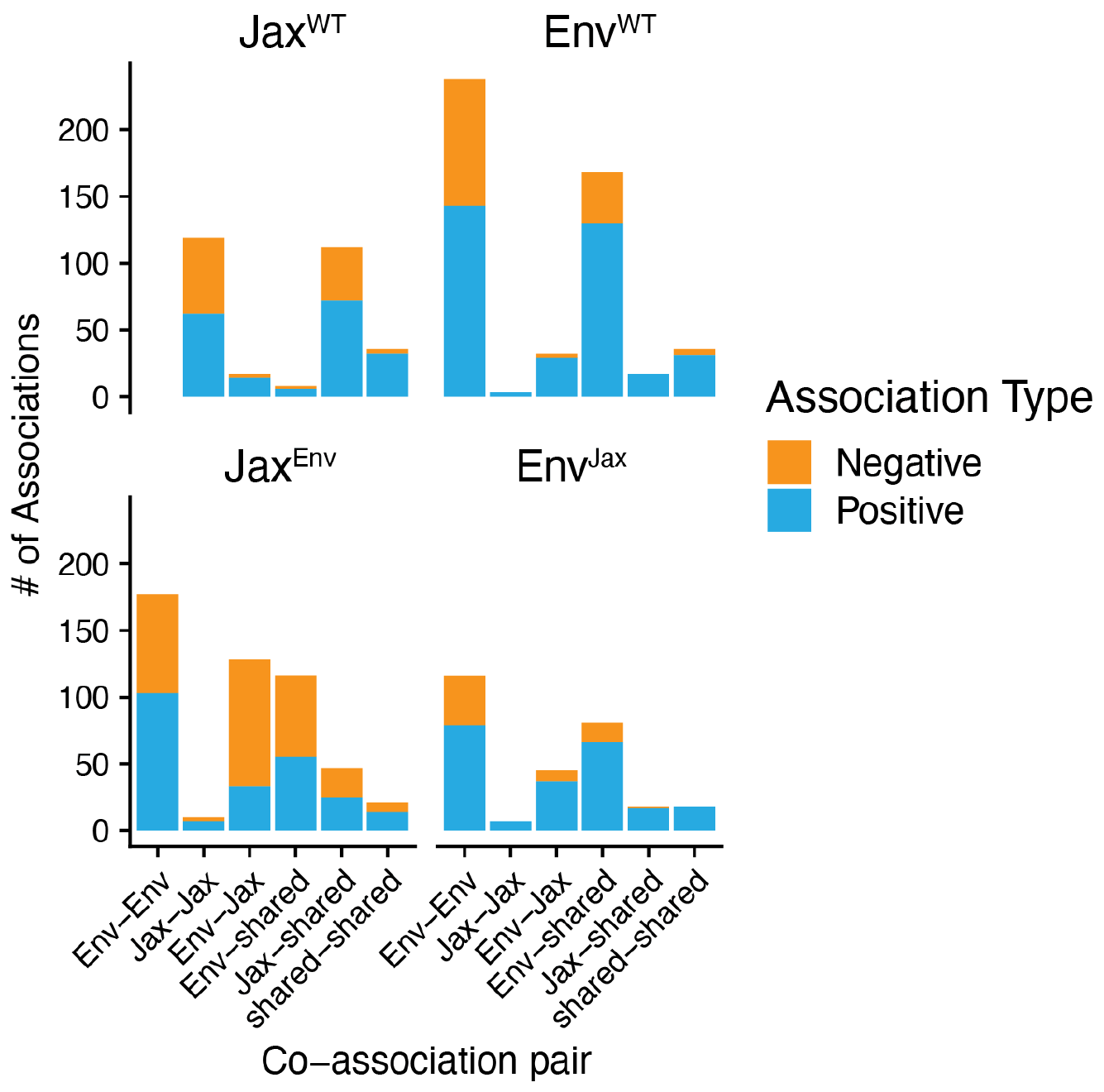
related to Figure 4. **Number of interactions between different types of microbial pairs**. Microbes were considered shared if they exhibited greater than 0.5% abundance in both Jax^WT^ and Env^WT^ populations whereas Env^WT^ and Jax^WT^ OTUs were only above this threshold in one of these groups. While Jax^WT^ contained many Jax-Jax spatial associations, relatively few were observed in Jax^Env^. Instead, Jax microbiota formed interactions with Env (Env-Jax) and Shared (Jax-shared) microbiota.

**Supplementary Figure 6.**
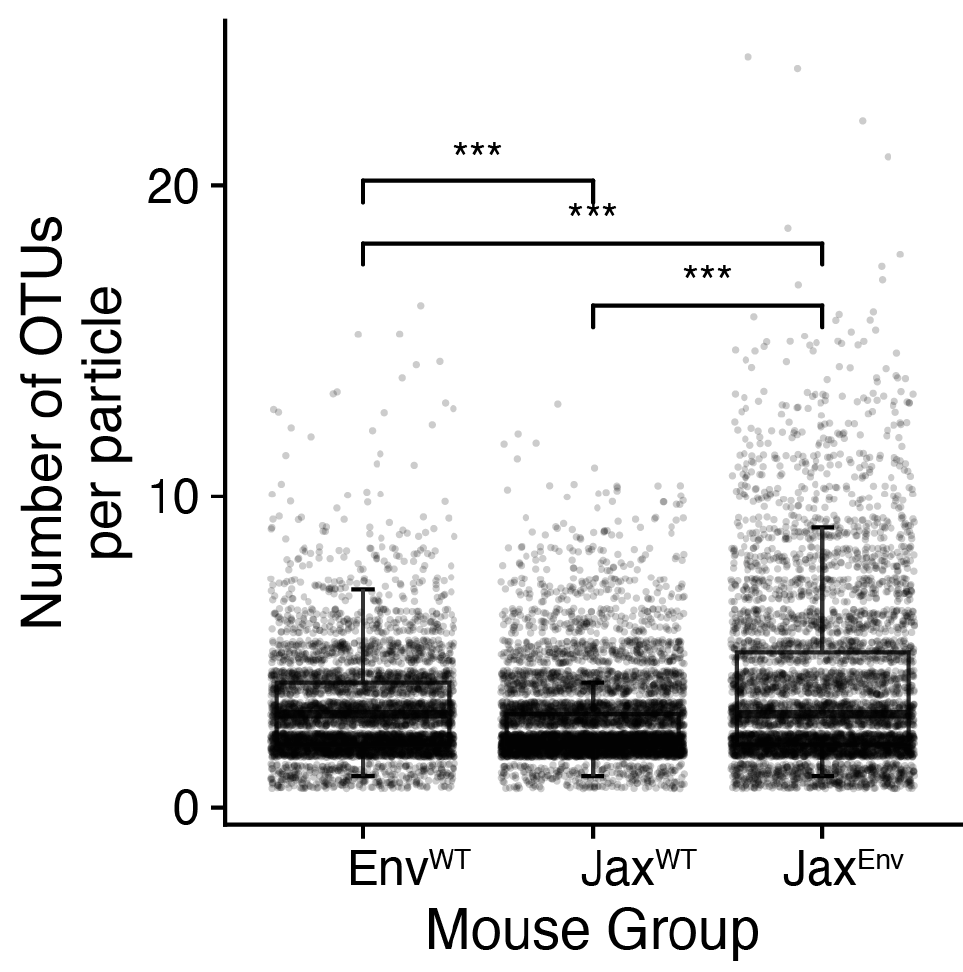
related to Figure 4. Number of distinct OTUs identified within particles derived from Env, Jax^Env^, and Jax^WT^ mice. Env mean: 3.01, Jax^Env^ mean: 3.76, Jax^WT^ mean: 2.88 (*** = *p* < 0.001, Wilcoxon rank-sum test).

**Supplementary Figure 7.**
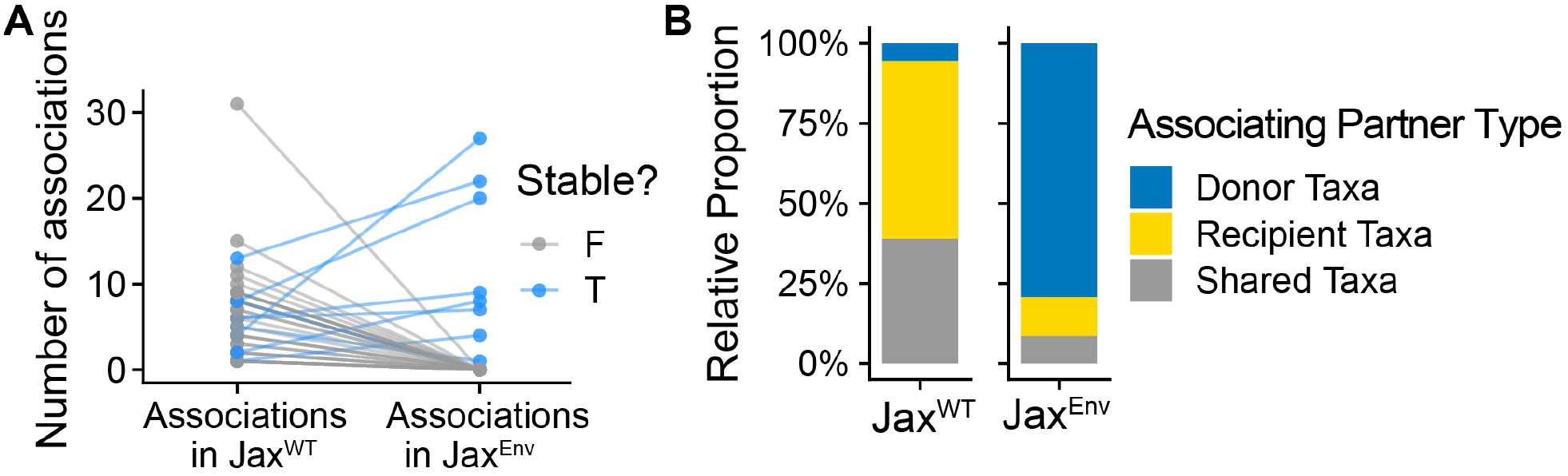
related to Figure 4. **Jax OTU interactions in native and donor microbiomes. (A)** Number of associations by Jax OTUs in Jax^WT^ mice and Jax^Env^ mice. Stable Jax microbiota show an increase in the number of associations they engage in post-FMT. **(B)** Composition of associating partners by Jax OTUs in Jax^WT^ mice and Jax^Env^ mice. Stable Jax microbiota establish spatial associations with donor taxa post-FMT.

**Supplementary Figure 8.**
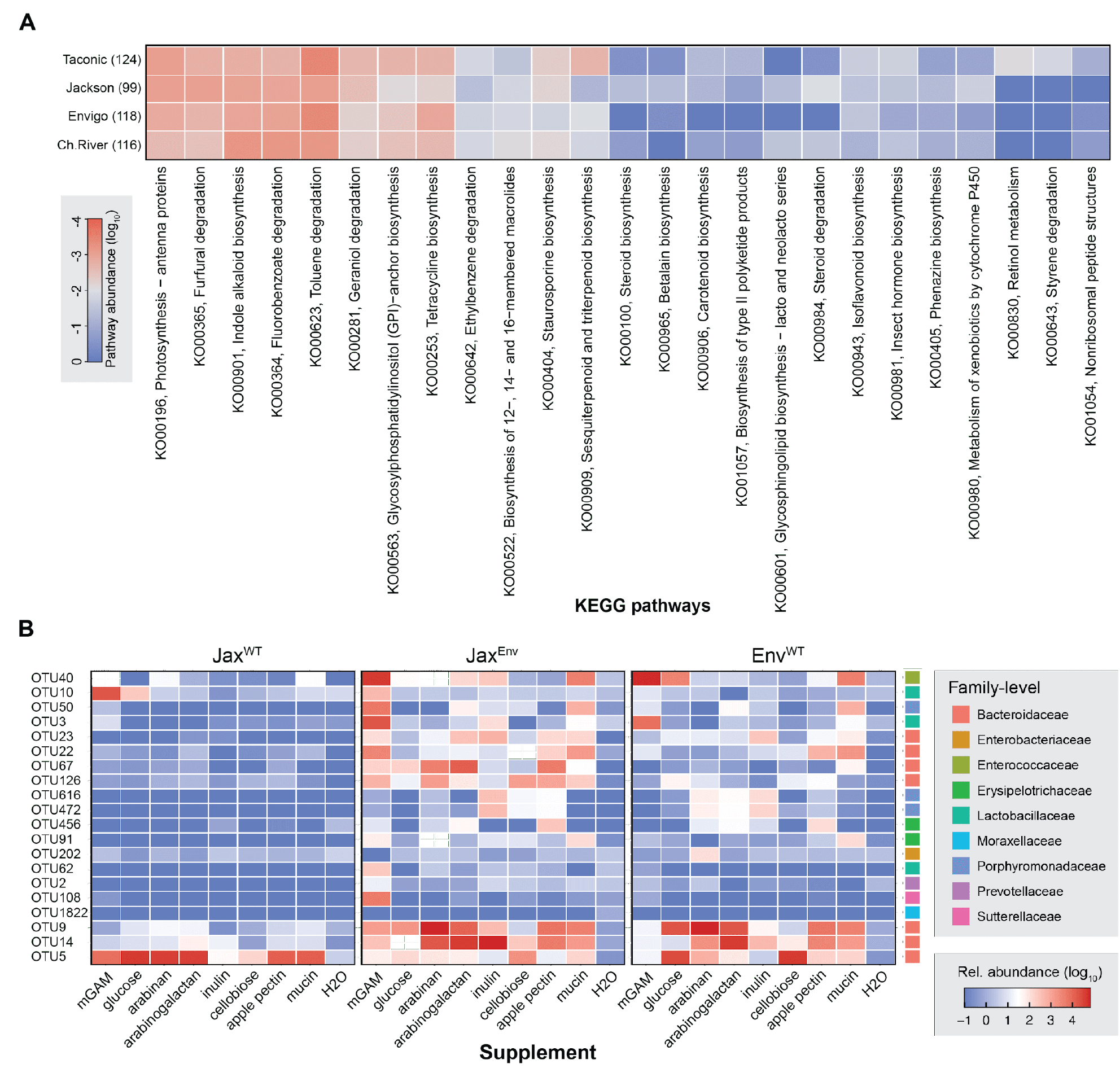
16S profiling of fecal microbiome communities after culturing. **(A)** Abundance of KEGG-annotated pathways identified amongst MAGs from each mouse vendor. Pathways presented exhibited the greatest variability between microbiomes. **(B)** Fecal communities were inoculated into *Bacteroides* minimal media supplemented with the indicated carbohydrate sources. Post FMT fecal pellets were harvested after five days of cohousing Jackson mice with Envigo.

**Supplementary Figure 9.**
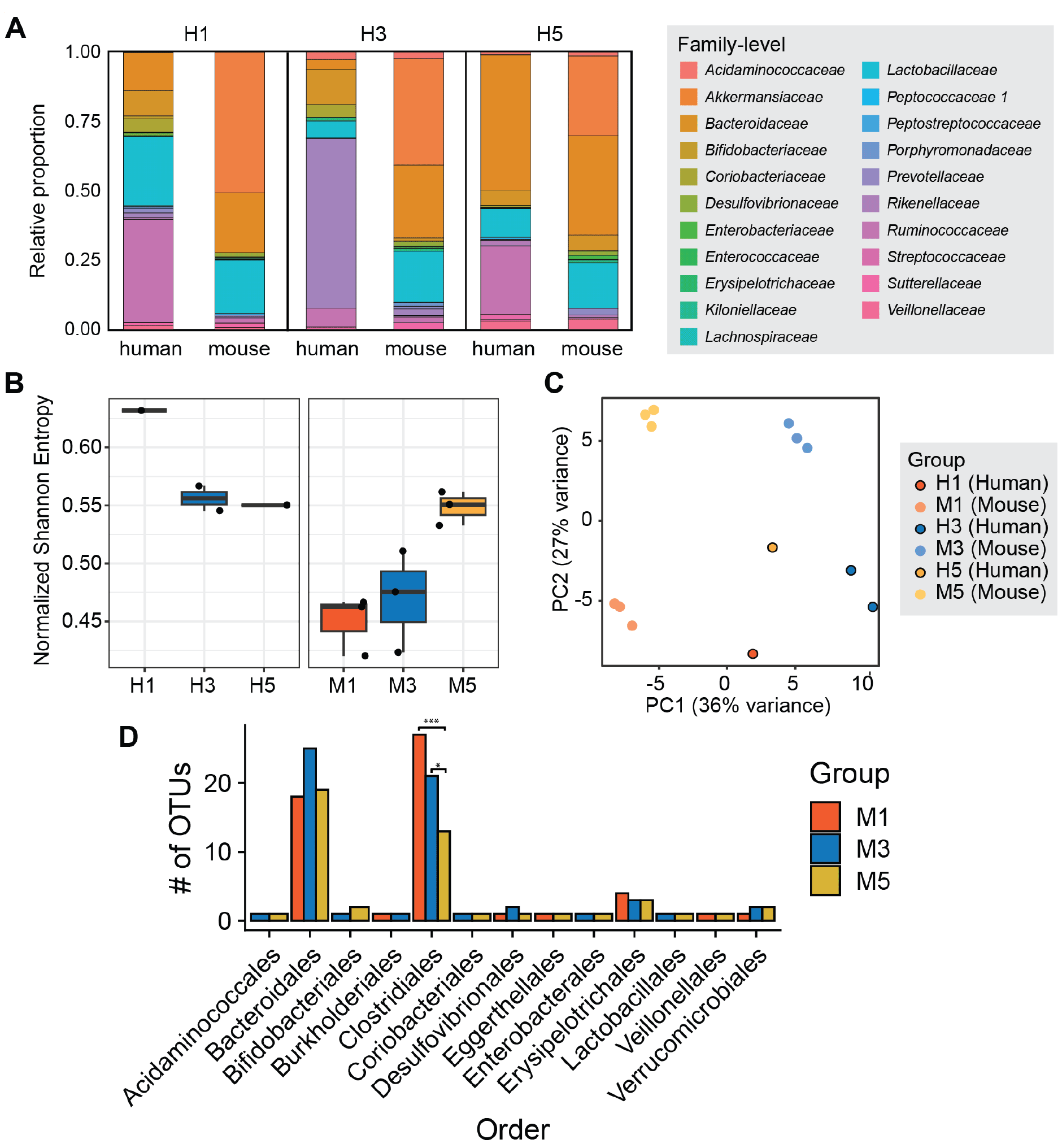
16S profiling of human donors and humanized mice. A) Family-level OTU composition of human fecal donors and corresponding humanized mice. Two sampling replicates were performed for human H3. **B)** Normalized Shannon Entropy of (left) human fecal donors and (right) humanized mice (N = 3 / donor). **C)** PCA of microbiome composition between human donors and humanized mouse cohorts at day 0 prior to mouse cohousing. **D)** Number of OTUs corresponding to each species in humanized mouse microbiomes. M1 and M3 mice contain significantly more *Clostridiales* OTUs than M5. (Wilcoxon rank sum test, M1-M5: *p* = 0.00074, M3-M5: *p* = 0.027).

